# Repeated application of bifocal transcranial alternating current stimulation (tACS) improves network connectivity and driving performance: a double-blind sham control study

**DOI:** 10.1101/2025.02.07.636997

**Authors:** Hakuei Fujiyama, Vanessa K. Bowden, Alexander D Tang, Jane Tan, Elisha Librizzi, Shayne Loft

## Abstract

Mounting evidence suggests that transcranial alternating current stimulation (tACS) can enhance response inhibition, a cognitive process crucial for sustained effort and decision-making. However, most prior studies have focused on within-session effects, with limited investigation into the effects of repeated applications, which are crucial for clinical applications. We examined the effects of repeated bifocal tACS targeting the right inferior frontal gyrus (rIFG) and pre-supplementary motor area (preSMA), regions implicated in response inhibition, on inhibitory control. We also explored changes to functional connectivity and whether this stimulation improved simulated driving performance. Thirty young adults (18–35 years) were assigned to either a sham or tACS group (20 Hz, 20 minutes), undergoing five stimulation sessions over two weeks. Resting-state electroencephalography (EEG) was used to assess functional connectivity between the preSMA and rIFG during the first and fifth bifocal tACS sessions and at a 7-day follow-up. Response inhibition was measured using a stop-signal task (SST) administered throughout the sessions. Participants completed two simulated driving tasks (braking, general driving) before the first and after the final tACS intervention. Results revealed a significant improvement in functional connectivity in the tACS group across sessions, although no changes were observed in response inhibition and the braking task. Notably, general driving performance improved, with participants demonstrating closer adherence to the speed limit and greater spare attentional capacity. These findings highlight the potential of repeated bifocal tACS to enhance functional connectivity and related cognitive and motor processes, suggesting promising clinical applications for addressing issues related to cortical connectivity.

---

Executive function is a set of higher-order cognitive processes that enable goal-directed behaviour, problem-solving, and adaptive responses to novel or complex situations (Blair, 2016). One of the key components of executive function is response inhibition, which plays a vital role in the continuous monitoring, adapting, and updating of behaviours in response to changes in the environment (Mostofsky & Simmonds, 2008). Impaired response inhibition has been linked to various psychological disorders characterised by impulsive behaviour, including addiction, attention deficit hyperactivity disorder, obsessive compulsiveness, and eating disorders (Bartholdy et al., 2016; Lipszyc & Schachar, 2010; Smith et al., 2014). Specifically, maladaptive response inhibition often manifests as slower execution of stopping responses or a diminished ability to delay the initiation of behaviours compared to healthy controls (Schmitt et al., 2018).

Response inhibition is also particularly important in everyday tasks requiring sustained effort and decision-making, such as driving, where safe driving often requires the inhibition of prepotent responses (Adrian et al., 2019; Hatfield et al., 2017), and where distractions can lead to catastrophic consequences (Beanland et al., 2013; World Health Organization, 2015). Considering the essential role of response inhibition in daily functioning and its identification as a core deficit in multiple psychological disorders, it is crucial to develop effective strategies to enhance response inhibition. Such advancements could pave the way for new intervention techniques that improve the quality of life and safety for those experiencing issues related to response inhibition (Bari & Robbins, 2013).

Understanding the neural mechanisms of response inhibition is crucial for improving its functionality. Key regions involved include the right inferior frontal gyrus (rIFG), pre-supplementary motor area (preSMA), and the subthalamic nucleus (STN), forming a cortical network essential for inhibitory control(e.g., Aron & Poldrack, 2006). Imaging studies have shown associations between successful inhibition in stop-signal tasks and increased activation in the rIFG and preSMA (e.g., Dambacher et al., 2014; Erika-Florence et al., 2014; Hampshire et al., 2010; Simmonds et al., 2008; Tsvetanov et al., 2018). Furthermore, transcranial magnetic stimulation (TMS) studies support the causal role of these regions, showing that disruption of rIFG or preSMA during stop trials impairs inhibitory control (Allen et al., 2018; Obeso et al., 2013).

Notably, a body of research suggests that the rIFG and preSMA work together to transmit neural signals through the basal ganglia to the primary motor cortex, effectively suppressing and cancelling motor response. Studies using functional magnetic resonance imaging (fMRI) have revealed extensive functional connections between the rIFG and pre-SMA during response inhibition, with both regions interacting with the basal ganglia (Jahfari et al., 2012). While some suggest only preSMA directly connects to the basal ganglia (Coxon et al., 2012; Duann et al., 2009; Rae et al., 2015), evidence supports robust connectivity between rIFG and preSMA, particularly during successful inhibition (Dambacher et al., 2014; Jahfari et al., 2012; Rae et al., 2015). TMS findings reveal that disrupting preSMA affects the influence of the IFG on motor output (Neubert et al., 2010) and enhancing preSMA activity strengthens connectivity within the fronto-basal-ganglia circuit (Xu et al., 2016). These results underscore the critical role of rIFG-preSMA connectivity in response inhibition, offering potential targets for enhancing inhibitory control.

One technique that has the potential to improve the functional connection between the rIFG and preSMA is transcranial alternating current stimulation (tACS)(Antal & Paulus, 2013). tACS is a non-invasive neuromodulation technique that applies weak oscillating electrical currents to the scalp. This technique is believed to affect the rhythmic activity of the brain by aligning endogenous neural oscillations with the frequency of the applied stimulation (Herrmann et al., 2013). When tACS is administered concurrently to two distinct cortical areas (i.e., bifocal tACS), it has the potential to enhance functional connectivity and communication between stimulated regions. Polanía and colleagues (2012) provided empirical evidence that bifocal tACS can modulate functional connectivity between cortical regions during a working memory task. Using 0° relative phase (in-phase) and 180° relative phase (anti-phase) stimulation over the left prefrontal and parietal cortices, they found that in-phase stimulation improved working memory performance, whereas anti-phase stimulation decreased performance. These findings suggested that tACS may influence the phase alignment of endogenous oscillations within fronto-parietal networks that are essential for working memory. For response inhibition, our previous study (Fujiyama et al., 2023) demonstrated that in-phase bifocal tACS at a beta frequency (20 Hz) over the rIFG and preSMA enhanced response inhibition and strengthened task-related functional connectivity, as observed with EEG. Similarly, other studies have reported favourable effects of bifocal tACS on a variety of cognitive and motor functions. (e.g., Grover et al., 2022; Lebihan et al., 2025; Meng et al., 2023; Reinhart & Nguyen, 2019; Violante et al., 2017).

While these findings highlight the potential of bifocal tACS to improve inhibitory function, most studies to date only examined its effect within a single session. An exception is a study conducted by Grover et al. (2022), who applied repeated bifocal tACS over four days and observed a positive impact on working memory, yet changes in functional connectivity were not considered in their study. It is, therefore, important to establish whether changes in functional connectivity and related improvements in cognitive function (behaviour) are retained over an extended period, which is a crucial attribute of any clinically meaningful intervention. Notably, no existing research has investigated whether multiple sessions of bifocal tACS over the rIFG and preSMA can result in long-lasting effects on functional connectivity and response inhibition.

The current study, for the first time, investigates whether bifocal tACS can induce lasting changes in functional connectivity between the rIFG and preSMA and improve response inhibition over an extended period of seven days. In addition, we explored the effects of repeated bifocal tACS application over the rIFG and preSMA on response inhibition. In addition to measuring the effect of stimulation on stop signal task (Congdon et al., 2012) performance in a laboratory setting, we extended our investigation to a real-world task requiring inhibition—simulated driving. This approach allowed us to evaluate the broader applicability of bifocal tACS and assess whether improvements in inhibitory control could extend to more complex, everyday tasks. We chose driving as it relies on executive functioning, including response inhibition (Adrian et al., 2019; Hatfield et al., 2017).

Specifically, drivers need to sustain attention and appropriately allocate sufficient cognitive resources to control their vehicle, navigate to a destination, and respond to hazards, which often requires inhibition of prepotent responses. Distraction, which occurs when attention is captured by other tasks, such as mind-wandering (Albert et al., 2022), looking at an engaging billboard (Crundall et al., 2006), or by a mobile phone (Gariazzo et al., 2018) can also lead to significant driving performance decrements. Since resisting distraction has been linked with response inhibition (Bissett et al., 2017), therefore enhancing response inhibition could potentially improve driving performance and safety. By investigating the effects of bifocal tACS on both lab-based response inhibition tasks and simulated driving, we aim to bridge the gap between controlled experimental settings and real-world applications. This approach allows us to assess the potential of tACS as a tool for improving cognitive functions critical for everyday activities, particularly those requiring sustained attention and inhibitory control.

## Method

### Participants

All potential participants underwent screening for contraindications to non-invasive brain stimulation and driving tasks, with exclusions made for individuals with psychiatric or neurological conditions or those taking medications incompatible with the stimulation protocol (Rossi et al., 2021). Handedness was assessed using the Edinburgh Handedness Inventory (Oldfield, 1971) and only participants who scored above 40, indicating right-handedness, were included in the study. This criterion was based on evidence linking left-handedness to variations in motor cortical representations and differing aftereffects of non-invasive brain stimulation (Fitzgerald et al., 2021; Nicolini et al., 2019). Prior to their participation in the study, all participants provided written informed consent. This study was approved by the Murdoch University Humans Ethics Committee (2020/186).

A power analysis to estimate the required sample size was performed using the “simr” package (Lenth, 2020) in R statistical package, version 4.4.1 (R Core Team, 2024), drawing on simulation data from our previous studies investigating the effect of non-invasive brain stimulation on behavioural measures (Fujiyama et al. 2016; 2022). With an alpha level set at 0.05 and a target power of 0.90, we estimated that 32 participants would be sufficient to detect a medium pre-post tACS change in response inhibition within a group (Cohen’s *d* = 0.4) based on a study that captured medium to large effect sizes on working and long-term memory task performances following multiple tACS sessions (Grover et al., 2022). Estimating an ∼ 20% attrition rate (Lansbergen et al., 2007), we recruited an additional 8 participants, resulting in a total of 40 participants.

Of the 40 participants recruited via the Murdoch University Research Participant Portal, four withdrew from the study due to scheduling conflicts, and one withdrew due to motion sickness after the first driving session. As a result, the current study started with 35 participants (Mean_age_ = 23.16, SD_age_ =4.89 yrs, 26 females). After the first tACS session, two participants dropped out due to scheduling conflicts. Two additional participants withdrew for the same reason after the third session, and one other participant did not return after the fourth session for undisclosed reasons. This resulted in 30 participants (Mean_age_ = 23.29, SD_age_ = 3.65 yrs, 21 females) completing all five tACS intervention sessions. For follow-up assessments, one participant in the tACS group did not complete the 7-day follow-up due to a scheduling conflict, resulting in 29 participants (Mean_age_ = 23.33, SD_age_ = 4.54 yrs, 20 females) completing all sessions. To address missing data, we did not use case-wise deletion. Instead, we included all available data at each time point in the analysis by employing a linear mixed-effects model, accommodating missing observations. Nevertheless, the results should be interpreted with some degree of caution due to the attrition rate and the reduced sample size at follow-up.

Participants were pseudo-randomly allocated to either the tACS or sham group to maintain balance as recruitment progressed. Allocation was adjusted iteratively to ensure an approximately equal number of participants in each group while accounting for ongoing recruitment dynamics. Consequently, both groups had 15 participants (10 female) with similar age profiles (tACS = 24.27 yrs, sham = 22.69 yrs, *p* = 0.53), noting that one of the participants in the tACS group did not participate in the 7-day follow-up.

### Materials

#### Stop Signal Task

Response inhibition was measured using a stop signal task (SST), a task that is considered a valid measure of latent response inhibition (Congdon et al., 2012). PsychoPy version 1.90.3 (Peirce et al., 2019) was used to program an 80-trial two-option reaction time SST program on a computer with a monitor refresh rate of 144Hz. As illustrated in Figure 1, each trial began with a white fixation cross presented in the centre of a black screen, which was then presented for 500ms. Subsequently, an imperative go signal (green arrow indicating left or right) was displayed, prompting the participant to press the corresponding arrow on the computer keyboard using their right index finger as fast as possible. Participants were instructed to start each trial by placing their index finger on the downward arrow key, which is situated between the left and right arrow keys, to avoid biased responses. Reaction time (RT) was measured between the onset of the go signal (green arrow) and the registration of the button press. Visual feedback was presented for all go trials, with RT presented in milliseconds on the screen after each successful go trial, “wrong response” for an incorrect response, and “missed” for no response. In 25% of the trials, the green Go signal was followed by a Stop signal (red arrow) facing the same direction as the green Go signal, which prompted participants to withhold their initiated response. The stop signal delay (SDD), the time between the onset of the Go signal and the stop signal, was manipulated in a staircase fashion (Verbruggen et al., 2019) across all blocks with an initial SSD of 200ms and a minimum SSD of 50ms. SSD increased by 50ms in the subsequent trial for every successful stop, while SSD decreased by 50ms in the subsequent trial for every failed stop signal response (Allen et al., 2018; Aron & Poldrack, 2006). Visual feedback was given, with “good!” presented for every successful stop and “try to stop” following every failed stop to encourage participants to perform their best.

**Figure 1.**
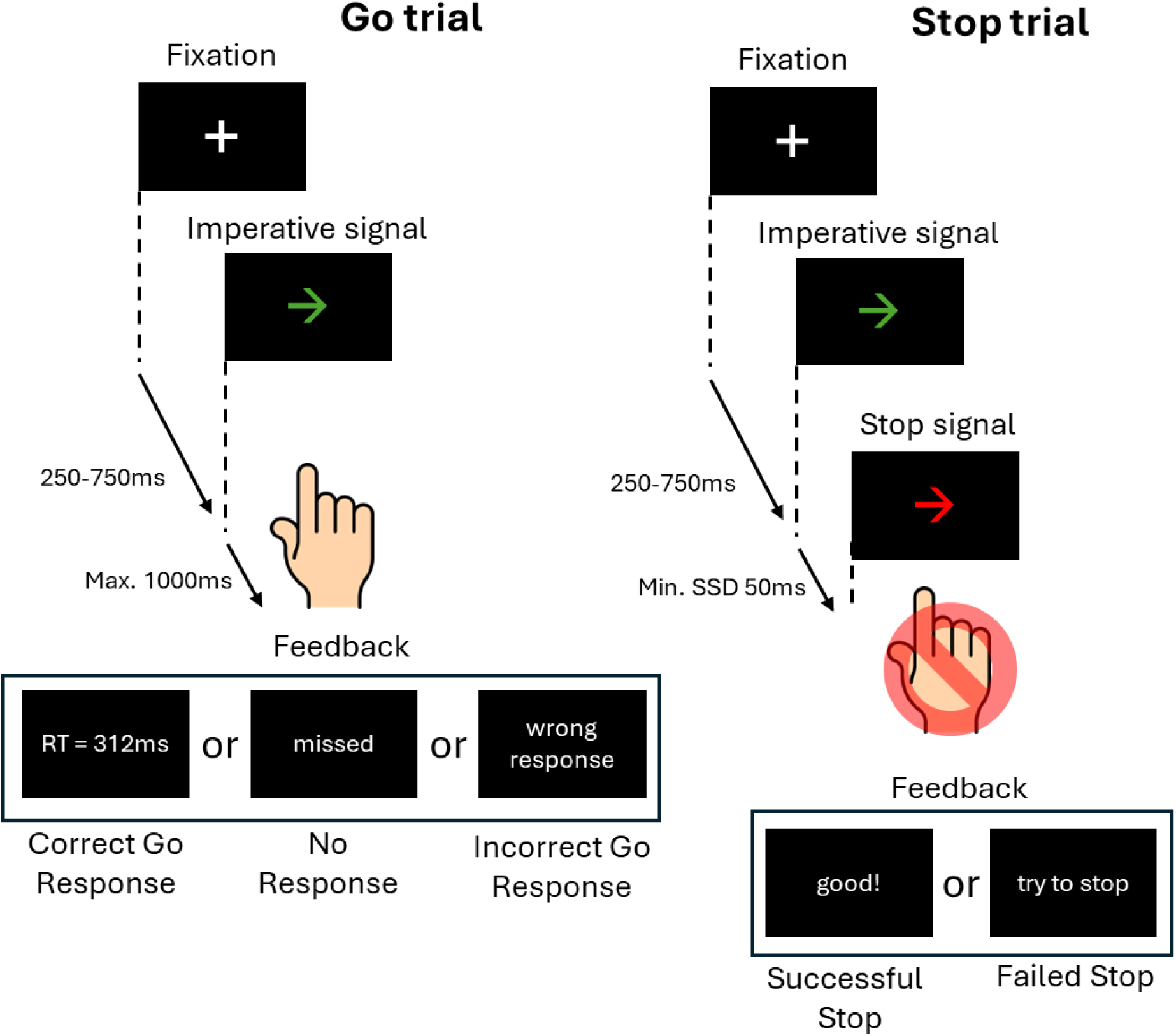
Schematic illustration of trial sequence in the stop signal task. Participants were instructed to respond to an imperative Go signal (green left- or right-pointing arrow) using their right index finger by pressing the corresponding key on a computer keyboard. Participants kept their index finger on the down arrow key on the keyboard between trials to avoid biasing the response. During stop trials (25% of the total trials), where the previously presented green arrow would turn red after a dynamic delay (stop signal delay, SSD), participants had to withhold their response. SSD on the subsequent stop trial was increased or decreased by 50ms following successful or unsuccessful stopping, respectively.

#### Driving simulator tasks

The medium-fidelity driving simulator used Oktal’s SCANeR Studio software (version 1.4) and featured three parallel 27-inch monitors mounted within an Obutto cockpit, providing a 135° wide-field display. Figure 3 shows the front windscreen view displayed on the central monitor, including a digital speedometer and a central rear-view mirror. Participants sat approximately 85cm away from the central monitor and operated a right-hand-drive vehicle using a Logitech steering wheel and pedal set. Vehicle speed and position data were recorded continuously at 1000 Hz and down-sampled to 50 Hz for analysis.

For the main driving task used to assess response inhibition, participants completed a 10-min braking task based on Muhrer and Vollrath (2011; also see Facchin et al., 2023). The braking task required participants to react as quickly as possible to an external event to change their routine driving behaviour. Participants were instructed to follow a lead car travelling at 50km/h on a straight section of road, while maintaining a safe following distance (Figure 2). The lead car was programmed to brake 8-11 times over a 3km distance. One lead car braked predictably (i.e., every 20s) before turning off the road. The participant then followed a second lead car which subsequently braked unpredictably (20-40s between braking events). The order of lead car presentation was counterbalanced.

**Figure 2.**
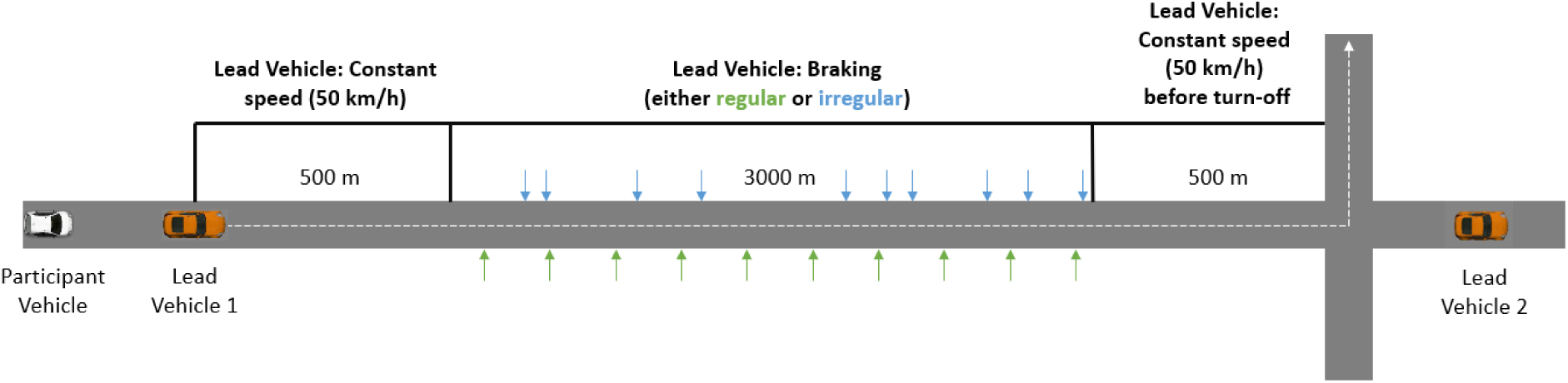
Braking task schematic. The participant followed a lead vehicle for 500m at constant speed before the lead vehicle started braking either regularly or irregularly. Once lead vehicle 1 turned off the main road, the participant encountered lead vehicle 2. The braking process was identical for lead vehicle 2, except for the pattern of braking was alternated (e.g., if lead vehicle 1 braked regularly, then lead vehicle 2 braked irregularly).

For generality, we also assessed fundamental vehicle control abilities. Participants drove in the inside-right lane of a continuous four-lane road, which was free of traffic, while the remaining lanes had light traffic (approximately five vehicles per minute). During a 5-minute practice scenario, participants were instructed to drive as close to the 50km/hr speed limit as possible while remaining in their lane. Participants then drove for ∼20 minutes.

While driving, participants also completed a visual Detection Response Task (DRT; Jahn et al., 2005). As illustrated in Figure 3, this task required them to respond to peripherally presented red dot targets (0.34° of visual angle) that appeared on the central monitor within an area 2° to 4° above the horizontal midline and 11° to 23° to the left of their forward viewpoint (Olsson & Burns, 2000; Patten et al., 2006). Targets remained visible for up to 2s, or until a response was made. Subsequent targets were separated by 6-16s. Participants responded to targets by pressing a button on the steering wheel with their right thumb, ensuring their hands stayed on the wheel (Bowden et al., 2017, 2019; Martens & Van Winsum, 2000). Participants’ speed and accuracy in responding to DRT targets can be considered as a measure of their spare cognitive capacity, thereby providing an immediate and quantifiable measure of the latent ‘cognitive workload’ (Castro et al., 2019; ISO DIS 17488, 2016) required by participants to maintain fundamental vehicle control.

**Figure 3.**
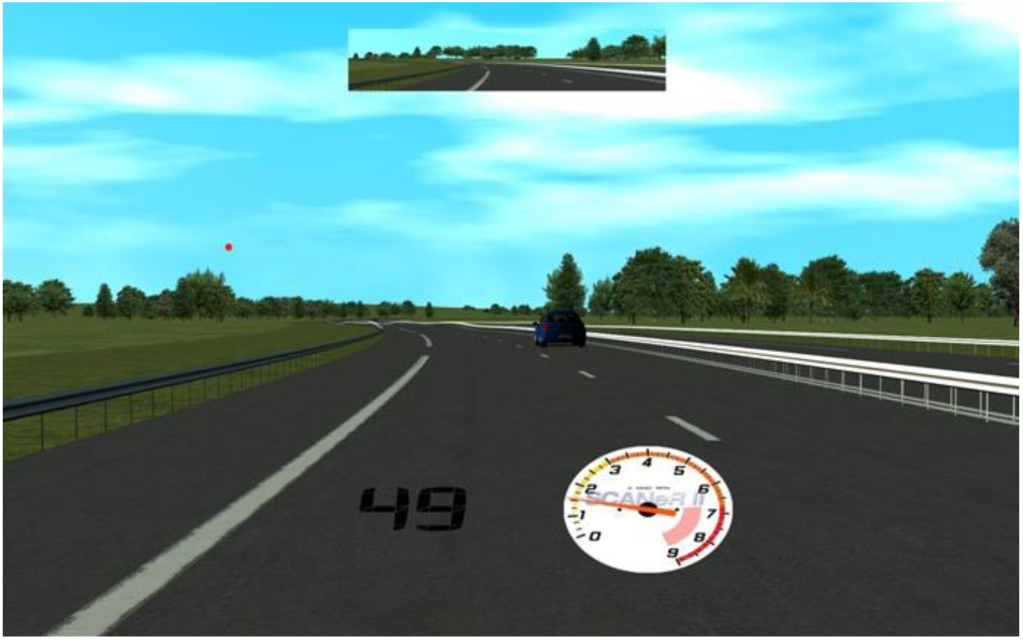
The central monitor view of the driving environment with rear-view mirror and digital speedometer displayed. A DRT target is presented above the horizon on the left.

#### Electroencephalography (EEG)

EEG was recorded using a 128-electrode EEG HydroCel Geodesic Sensor Net (Magstim EGI, Eugene, OR). Net Station (5.4.2) software recorded sensor-level EEG signals from Ag-AgCl scalp electrodes. Signals were amplified using a Net Amps 300 amplifier, low and high pass filtered (0.1-500 Hz) with a 1000 Hz sample rate. Electrode impedance was kept below 50 kΩ as recommended by the manufacturer (Magstim EGI, Eugene, OR). During resting-state EEG recording, participants viewed a fixation cross presented on a PC monitor for 3 minutes.

#### Transcranial Alternating Current Stimulation (tACS)

Bifocal tACS was administered using the neuroConn DC-STIMULATOR MC machine (NeuroConn, Ilmenau, Germany). Three rubber electrodes of 2 cm in diameter were applied to the scalp using Ten20 conductive paste in a 2 x 1 montage (Figure 4), targeting the rIFG and preSMA. The electrode placement was determined using the international 10-20 system. The preSMA was located at Fz and the rIFG in the intersection between Fz to T4 and Cz to F8. Alternating currents were applied using a beta-frequency of 20 Hz. A current intensity (peak-to-peak amplitude) of 1 mA was applied to the rIFG and 1.6 mA to the preSMA with zero DC offset, which was informed by the e-field modelling presented elsewhere (Tan et al., 2020).

**Figure 4.**
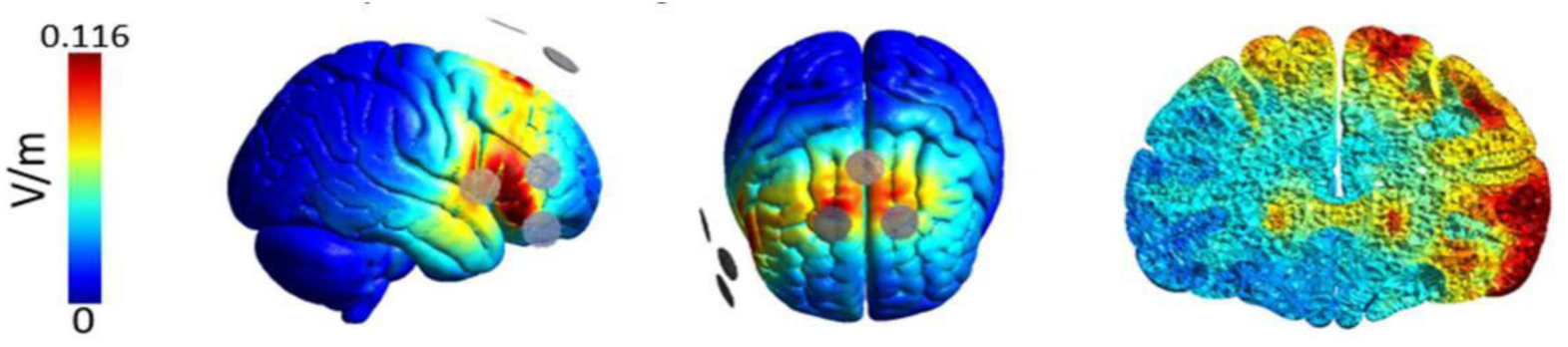
Electrical current flow modelling for the tACS (2 x 1) electrode montage over the right inferior frontal gyrus and pre-supplementary motor area (Adapted from *The importance of model-driven approaches to set stimulation intensity for multi-channel transcranial alternating current stimulation (tACS),* by Tan et al., published in *Brain Stimulation*, 2019, under a CC BY 4.0 license. Vol75 (mesh volume with field strength ≥ 75% of the 99.9th percentile) and Vol50 (mesh volume with field strength ≥ 50% of the 99.9th percentile) provide measures of focality. The grey circles represent the rubber electrodes (2 cm diameter) with two active electrodes placed anteriorly and one return electrode placed poster to the target region. The current amplitude (1mA peak-to-peak) was equally split between the two active electrodes.

The currents were delivered using a 0° phase to the preSMA and rIFG, indicating that the peaks and troughs of the electrical waves were coupled in both regions. The stimulation was sustained for 20 minutes, a period chosen to allow sufficient time for the neural circuits to respond to the electrical modulation (De Koninck et al., 2023). In the sham condition, the tACS stimulation ramped up to the respective current intensity and immediately ramped down at the beginning and end of the 20 minutes for 30 seconds to simulate stimulation (Woods et al., 2016). The tACS machine was pre-programmed by a research associate with codes representing the two conditions to ensure both participant and researcher blinding.

#### Control Measures

##### Sleep, Caffeine, and Alcohol Questionnaire (SCA-Q)

The Sleep, Caffeine and Alcohol Questionnaire (e.g., Fujiyama et al., 2023) was used to account for potential confounding factors affecting the response to tACS, such as sleep quality, hours slept, and caffeine and alcohol intake within the 12 hours prior to the session. Sleep quality was rated on a scale from 1 (poor) to 10 (excellent). Participants reported caffeine consumption in milligrams and alcohol intake in standard drinks, with reference tables provided to help accurately estimate the caffeine content of various beverages and the standard drink equivalents of alcoholic beverages.

##### Transcranial Electrical Stimulation (tES) Questionnaire

Upon completion of each tACS session, a tES questionnaire (Fujiyama et al., 2022; 2023; Vancleef et al., 2016) was given to assess blinding adequacy and stimulation effects. The questions involved the sensations experienced and the side effects of the stimulation. The questionnaire includes 12 items of sensations that are commonly associated with non-invasive brain stimulation, such as “heat”, “headache”, and “tingling” (Woods et al., 2016). The items were rated on a five-point Likert scale, ranging from “nothing” to “very strong”. The same rating scale was used to ask participants whether the sensations affected their performance of the SST task, ranging from “a little” to “very much”. Furthermore, three items were asked about the time the sensations occurred during the stimulation with a three-point Likert scale, comprising of “at the start”, “in the middle”, and “at the end” of the stimulation.

### Procedure

The project timeline for each participant was consistent across both active and sham groups (Figure 5). Participants completed five experimental sessions of either active or sham tACS, followed by a 7-day post-intervention follow-up session. Resting-state EEG (rsEEG) was conducted during the first (baseline) and fifth (post-intervention) sessions. During rsEEG recording, participants were instructed to maintain a neutral gaze at a fixation cross for 3 minutes (eyes open), avoiding complex thought processes. Following the rsEEG, participants completed the SST during EEG recording. This began with a 16-trial practice session, followed by two blocks of 80 trials (pre-stimulation). tACS electrodes were then applied using conductive paste, and stimulation was administered for 20 minutes, including a 30-second ramp-up and ramp-down. During stimulation, participants completed three additional blocks of 80 SST trials (mid-stimulation). After the tACS session, another rsEEG recording was conducted, followed by two additional blocks of SST trials. The EEG nets remained in place during the tACS application, as the tACS electrodes were applied without needing to remove the nets. Subsequent experimental sessions (sessions 2–5) consisted only of the 20-minute tACS application and three blocks of 80 SST trials. While both resting-state and task-related EEG data were collected, this study focuses on rsEEG to examine baseline neural oscillatory activity. This aligns with the primary objective of exploring the effects of repeated tACS application on behaviour and functional connectivity.

**Figure 5.**
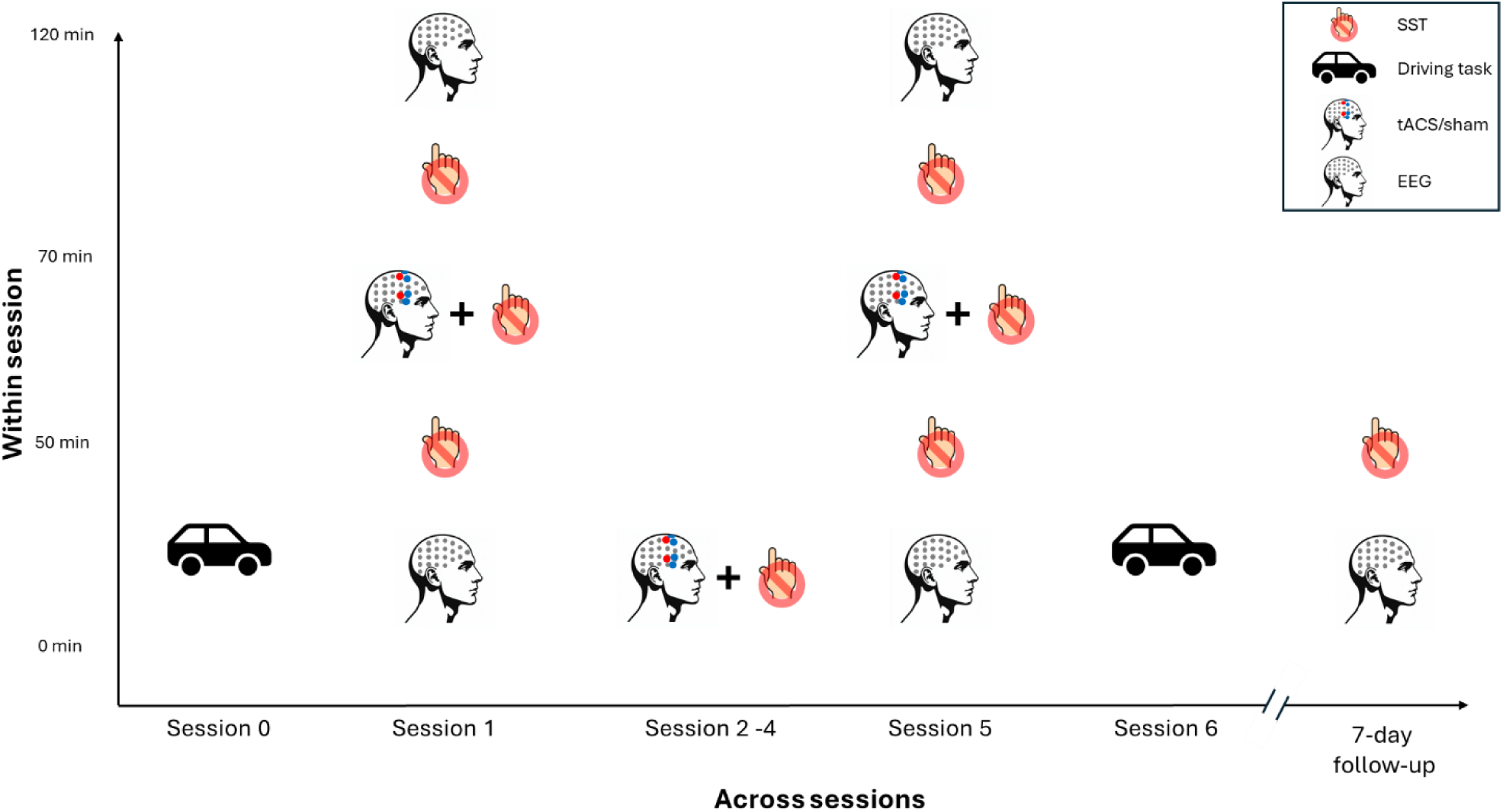
Project timeline within and across sessions. Participants received tACS over five sessions (Sessions 1-5) across a two-week period, with 1-2 days between each session. Resting-state EEG (rsEEG) was recorded in Sessions 1 and 5, as well as during follow-up sessions at 7 days post-intervention. Driving task performance was assessed in Session 0 (baseline) and Session 6 (post-intervention). In Sessions 1-5, participants performed the Stop Signal Task (SST) during tACS application. Additionally, in Sessions 1 and 5, the rsEEG and SST were administered both before and after tACS to examine within-session effects. During the follow-up sessions, the SST was performed without tACS.

The 7-day follow-up session included EEG recordings and two blocks of the SST. The first and fifth sessions lasted approximately 90 minutes, while the remaining sessions were around 45 minutes each. Driving performance was assessed at Session 0 (prior to the tACS intervention) and 1 day after the final tACS session (Session 6). The decision to separate the post-intervention driving assessment from the EEG and SSRT assessments was primarily due to logistical constraints. The tACS, EEG, and SST assessments were conducted at the Murdoch University campus, while the driving task was administered at The University of Western Australia campus. Given the need for specialised facilities and equipment at each location, it was impractical to perform all assessments at the same site.

### Data Processing and Statistical Analysis

#### EEG Pre-processing

EEG data were pre-processed using the EEGLAB toolbox RELAX (Bailey et al., 2023) through the MATLAB environment (MathWorks, R2023b). RELAX is a data-cleaning pipeline that reduces vascular, ocular, and myogenic-induced artifacts to preserve neural signal readings. Using RELAX, the data was down-sampled at 500 Hz. Then, a 50 Hz notch filter and a bandpass filter (1-80 Hz) were applied. Second, problematic EEG sensors and noisy time periods were removed. Third, eye blink and muscle movement-induced artifacts were removed using the Multiple Wiener Filtering method. The Wavelet Enhance Independent Component Analysis removed artifacts not detected in the previous method.

Last, once the removed EEG sensors and artifacts were interpolated, the data was divided into two-second segments for statistical analysis.

Functional connectivity between the rIFG and preSMA regions was measured using the imaginary coherence (ImCoh), which measures the consistency of phase differences between two signals (Nolte et al., 2004). Unlike the real part of coherency, the imaginary component is insensitive to artifacts caused by volume conduction (Nolte et al., 2004). Larger ImCoh values represent stronger phase synchrony between the two cortical areas, indicating the presence of a specific temporal relationship (Cattai et al., 2021). For ImCoh calculation, the time series of each electrode were convolved with complex Morlet wavelets for frequencies between 4 and 90 Hz in 1 Hz increments (87 wavelet frequencies in total). The length of the wavelets started at 3 cycles for the lowest frequency and logarithmically increased as the frequencies increased, such that the length was 13 cycles for the highest frequency. This approach balances temporal and frequency precision (Cohen, 1988). To minimise the effects of edge artifacts, analytic signals were obtained from time windows of 400 to 1600 ms (at 20 ms intervals) within each 2000 ms epoch. ImCoh values at 20 Hz were computed for each epoch using an electrode cluster centred on Fz and AFz to represent the preSMA, and a cluster consisting of FC6, F6, and F4 to represent the rIFG using the following formula:

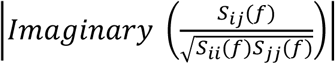

Here, *i* and *j* represented the time series of each electrode. For frequency *f*, the cross-spectral density *S*_*ij*_(*f*) was taken from the complex conjugation of the complex Fourier transforms *x*_*i*_(*f*) and *x*_*j*_(*f*). Coherency was extracted by normalising the cross-spectral density by the square root of the signals’ spectral power, *S*_*ii*_(*f*) and *S*_*jj*_(*f*). ImCoh values range from 0 to 1, where a value of 0 indicates completely random phase angle differences between the signals, while a value of 1 signifies perfect phase coupling between them.

#### SST calculation

The integration method (Verbruggen et al., 2019) was used to calculate stop signal reaction Time (SSRT), which served as a measure of response inhibition (i.e., action cancellation when the stop signal appears). The integration method is less affected by stopping success rates that deviate from 50% (Band et al., 2003). Specifically, for each participant and at each time point, the mean SSD was individually computed. In the initial stages of calculating the SSRT, all go trials were considered. This comprehensive approach included go trials where choice errors occurred, as well as those with premature responses, ensuring a thorough representation of the participant’s performance across various trial types (Verbruggen et al., 2019). Since SSRT estimation relies on the assumption of an independent race between the go and stop processes (e.g., Verbruggen & Logan, 2009), for each participant and each assessment time point, we excluded conditions if the mean RT on unsuccessful stop trials was greater than the mean Go RT in that same condition (Verbruggen et al., 2019). Furthermore, for each participant and for each time point, conditions with stop accuracy < 0.25 or > 0.75 were excluded, as suggested by Congdon and colleagues (2012) and Verbruggen et al. (2019). Go RT for correct responses in the corresponding trials was arranged in ascending order, and the Go RT corresponding to the stop success rate (i.e., 1-percent successful inhibition) was identified. For example, if the stop success was 55%, the 55^th^ percentile Go reaction time was identified as the quantile reaction time, and the mean SSD was subtracted from the quantile reaction time to estimate SSRT. We also excluded SSRTs less than 50 ms from the statistical analyses (Congdon et al., 2012). For Go RT analysis (but not for the calculations of SSRT), any trials with less than 150 ms of Go RT were excluded as they were deemed to be too fast (Perquin et al., 2024).

#### Driving task data processing

For the braking task, performance was only recorded during the braking interval (Figure 2), which excludes the constant speed interval in which participants accelerated to match the lead vehicle speed. We recorded speed variability (km/h) and time headway (THW) between the participant car and the lead car THW = separation / speed). THW during braking events indicates how participants adapted their behaviour to the lead car braking, with a larger headway allowing them to react more safely to braking events (Muhrer & Volrath, 2011).

For the general driving task, the first 45 seconds of driving were excluded to allow participants to accelerate. We then recorded speed and lane position variability (lower variability reflects improved speed/lane maintenance), median speed (km/hr), time spent speeding (>5km/hr over the limit). For the DRT, a hit was recorded if participants responded within 2.5s of target onset, and a false alarm for responses made outside this window. Median response time to hits is reported.

#### Statistical analysis

A generalised linear mixed model (GLMM) was constructed to analyse ImCoh and dependent variables for the driving task, while a linear mixed model (LMM) was employed for SSRT. For ImCoh and SSRT, two models were constructed. First, we constructed a model that analysed the within-session effect of tACS. For this model, the fixed factors of GROUP (tACS, sham), TIME (pre, post), and SESSION (session 1, session 5) were included for ImCoh and for SSRT an additional level in TIME (pre, during, post) was included in the model. Second, a model that analysed between-session effects examining the effect of repeated tACS application. For this model, the fixed factors of GROUP (tACS, sham) and SESSION (session 1, session 5,7-day follow-up [FU]) for ImCoh and an additional three levels in SESSION (session 1, session 2, session 3, session 4, session 5,7-day FU) for SSRT were included in the model using pre-stimulation data for each session, while for the driving tasks, the fixed factor SESSION had only two levels: Session 0 and session 6. Within-session changes in SST and ImCoh data from sessions 1 and 5 were analysed to evaluate the effects of tACS. For SST, the model included fixed factors of GROUP (tACS, sham) and TIME (pre, during, post). For ImCoh, the model used the same fixed factors, but TIME included only two levels (pre, post). For all models, the by-subject intercept was included as a random effect.

The assumptions for the models, including linearity, homogeneity of variance, and normal distribution of the model’s residuals, were evaluated using the “DHARMa” package (Hartig, 2024), which employs a simulation-based method to examine residuals for fitted G/LMMs. Null hypothesis significance testing for main effects and interactions was conducted with Wald Chi-Squared tests for the GLMM analyses and *F-*tests for LMM analyses, and significant main effects and interactions were further investigated with Bonferroni corrected contrasts where appropriate. To avoid potential misinterpretations and oversimplifications of the outcomes (Pek & Flora, 2018), standardised effect sizes for each fixed factor were not reported for the GLMM. To facilitate the interpretation, Cohen’s *d* for follow-up contrasts were provided alongside significance levels. Mann-Whitney tests were used to assess differences between groups in sleep duration, sleep quality, and caffeine and alcohol consumption at each session, while Wilcoxon’s signed-rank tests were used to assess differences between sessions for each group separately. For both Mann-Whiteney and Wilcoxon’s signed-rank tests, *r* value was provided as a measure of effect size for each comparison. Statistical significance for all tests was set at .05. In presenting results, the data were expressed as mean ± 95% confidence intervals, unless specified.

Statistical analyses and visual illustrations were performed using R statistical package, version 4.4.1 (R Core Team, 2024) with an integrated environment, RStudio version2024.04.2+764 (RStudio Team, 2015) for Windows using “lmerTest” v3.1-3 (Kuznetsova et al., 2020) to fit LMM and GLMM, “DHARMa” package (Hartig, 2024) for LMM assumptions of linearity, homogeneity of variance and normal distribution of residuals “emmeans” v1.10.3 (Lenth, 2024) for follow-up contrasts, “ggplot2”3.5.1 package (Wickham et al., 2016) for graphical plots, “dplyr” v1.1.4 (Wickham et al., 2023) for data manipulation, “janitor” v2.2.0 (Firke, 2023) for cleaning data, “here” v1.01 (Muller & Bryan, 2020) for declaring paths, “knitr” v1.47 (Xie, 2024) for report generation, “reader” v1.0.6 (Cooper, 2017) for reading data files, “reshape2” v0.8.1 (Wickham, 2007) for reshaping data, “skimr” v2.1.5 (Waring et al., 2022) for data summaries, “stringr” v1.5.1 (Wickham, 2023) for string operation wrappers, and ‘coin’v1.4-3 (Hothorn et al., 2023). All data and codes are publicly available on the Open Science Framework (OSF): https://osf.io/bdy4q/.

## Results

### Control Measures

Descriptive and inferential statistics for SCA and tES sensation ratings are shown in Table 1. Mann-Whitney tests revealed that there were no statistical differences between the tACS and sham group in sleep quality and duration, consumption of caffeine or alcohol, or sensation related to tACS in any sessions, suggesting a limited impact of the confounding factors on outcome measures and that participants were likely blinded from the conditions.

**Table 1.** Comparisons between tACS and sham group on sleep quality and duration, consumption of caffeine and alcohol, and sensation related to tACS.

|  | tACS |  | Sham |  | *p* | *r* |
| --- | --- | --- | --- | --- | --- | --- |
|  | *M* | *SD* | *M* | *SD* |  |  |
| Sleep quality (0-10) | 6.93 | 1.62 | 7.15 | 1.35 | >.31 | <\|.23\| |
| Sleep duration (hrs) | 7.47 | 1.23 | 7.12 | 1.24 | >.38 | <\|.22\| |
| Caffeine intake (units) | 37.55 | 38.95 | 29.53 | 52.62 | >.26 | <\|.25\| |
| Alcohol intake (units) | 0.46 | 1.54 | 0 | 0 | >.09 | <\|.20\| |
| tES sensation (0-62) | 10.03 | 5.39 | 7.93 | 5.48 | >.17 | <\|.28\| |

*Note:* descriptive statistics are average across sessions for each group. The smallest *p*-value and the largest absolute *r* value across tests that compared group differences at each session are presented. tES = transcranial electrical stimulation

Similarly, for each group, Wilcoxon’s signed-rank tests revealed no statistically significant differences between sessions on sleep quality and duration, caffeine and alcohol intake, and sensations related to the stimulation, *p*s > .42, |*r*s|<. 28. This indicated that sleep quality and duration, consumption of caffeine and alcohol, and sensation related to the stimulation were comparable across sessions within groups.

### EEG functional connectivity (imaginary coherence, ImCoh)

#### Within-session changes

None of the main effects and interactions was statistically significant, *χ^2^* < 0.69*, ps>* .42, suggesting that ImCoh did not change within sessions 1 or 5 for either group.

#### Between-session effect

There was a significant main effect of SESSION, *χ^2^* (1) = 45.66, *p* < .001, and interaction between GROUP and SESSION, *χ^2^*(2) = 139.89, *p* < .001. As shown in Figure 6, ImCoh in the tACS group showed a significant increase from Session 1 to Session 5, rising by 5.4%, *z* = 5.58, *p* < .001, *d* = 0.69. This improvement was maintained at the 7-day FU, where ImCoh was 3.9% greater than at Session 1, *z* = 3.52, *p* = .001, *d* = 0.48. In contrast, ImCoh in the sham group significantly declined from Session 1 to Session 5 by 6.9%, *z* = −5.74, *p* <.001, *d* = 0.83, and a further 7.4% reduction from Session 5 was observed at 7-day FU, *z* = −6.12, *p* <.001, *d* = 0.96. The group differences were evident across all three sessions; at Session 1, the tACS group had a significantly lower ImCoh value than the sham group, *z* = −2.67, *p* = .008, *d* = 0.79, while at Session 5 and 7-day FU, the tACS group had greater ImCoh values compared to the sham group, *z*s > 2.39, *p*s < 0.017, *d*s > 0.74.

**Figure 6.**
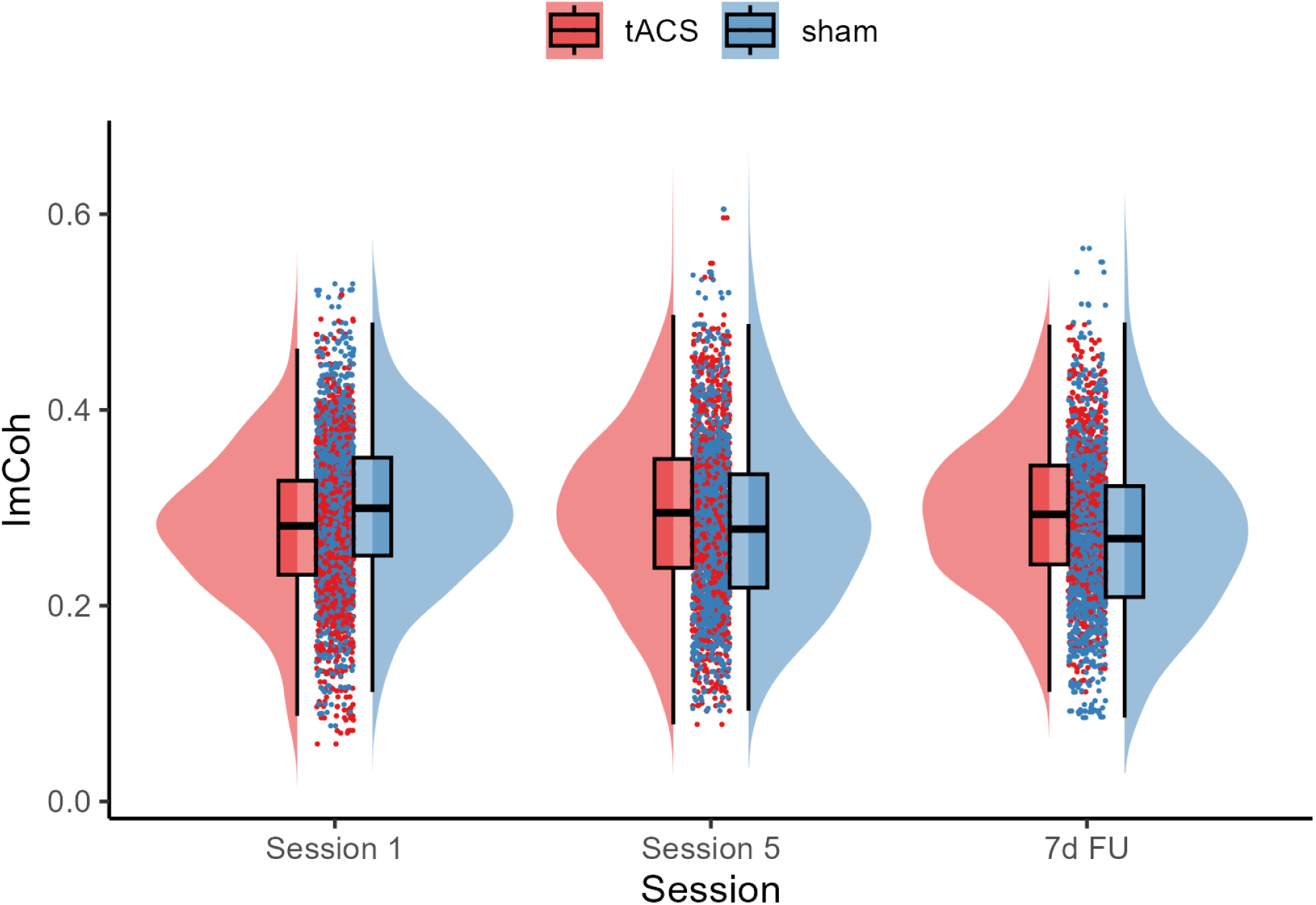
Imaginary coherence (ImCoh) changes across sessions for the tACS and sham groups. The boxplots show the median and interquartile range to visually summarise central tendency and dispersion. The horizontal line within the box plot represents the median. The top and the bottom lines represent the upper and lower quartiles, respectively. The data points outside of the whiskers are >1.5 quartiles.

### Stopping ability (stop signal reaction time)

#### Within-session changes

Contrary to the expectation, neither GROUP x TIME nor GROUP x TIME x SESSION interactions were statistically significant, *F*s < 0.48, *p*s > 0.61, suggesting that tACS did not have an impact on SSRT within Session 1 or 5. The only significant effect was the main effect of SESSION, *F* (1, 360.43) = 9.28, *p* = .003, indicating that, overall, SSRT was significantly faster in Session 5 (205.79 ± 4.01 ms) than in Session 1 (222.05 ± 3.64 ms). No other main effects and interactions were significant, *F*s < 2.60, *p*s > 0.08.

#### Between session changes

We observed a significant main effect of SESSION, *F*(5, 545.27) = 3.32, *p* = .006. As illustrated in Figure 7, the follow-up contrast showed that SSRT at 7-day FU (216.25 ± 7.51 ms) was significantly faster than Sessions 2 (228.44 ± 7.80 ms) and 4 (227.38 ± 7.93 ms), *t*s > 3.01, *p*s < .042, *d*s > 0.47. Since the interaction between GROUP and SESSION was not statistically significant, *F*(5, 545.27) = 1.18, *p* = .41, the observed improvement in SSRT at 7-day FU was unlikely due to the repeated application of tACS, but likely due to a generalised training effect.

**Figure 7.**
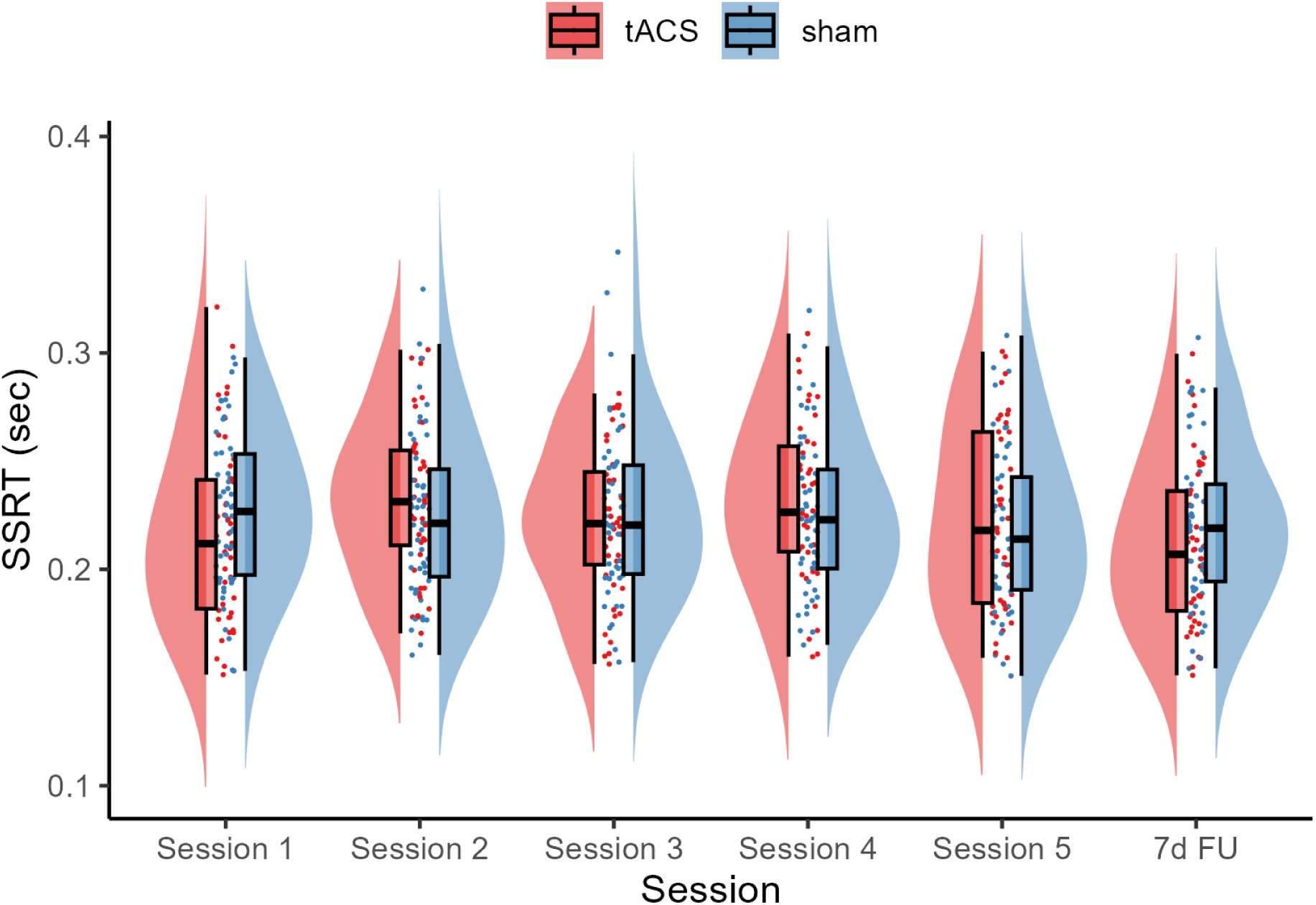
SSRT changes across sessions for the tACS and sham groups. The boxplots show the median and interquartile range to visually summarise central tendency and dispersion. The horizontal line within the box plot represents the median. The top and the bottom lines represent the upper and lower quartiles, respectively. The data points outside of the whiskers are >1.5 quartiles.

### Driving performance

Table 2 provides a summary of descriptive and inferential statistics for the braking task. There was no significant interaction between SESSION and TIME for any of the measurements, *χ^2^*s < 1.79, *p*s > 0.18, which suggests that tACS did not impact participants’ ability to maintain a safe following distance during this task.

**Table 2.** Descriptive and inferential statistics for **braking task** performance by group and time point. Time headway (THW) is in seconds. Regular refers to performance while following the regularly braking lead vehicle, while irregular is following the irregularly braking lead vehicle.

| Dependent Variable | Mean±CI |  |  |  | SESSION x TIME interaction<br>Chi-square value and p-value |
| --- | --- | --- | --- | --- | --- |
|  | tACS |  | sham |  |  |
|  | Session 0 | Session 6 | Session 0 | Session 6 |  |
| Speed variability (km/h) – Irregular | 0.37±0.19 | 0.34±0.18 | 0.21±0.07 | 0.33±0.41 | $\chi^2=1.79, p=.181$ |
| Speed variability (km/h) – Regular | 0.23±0.15 | 0.24±0.13 | 0.23±0.07 | 0.19±0.09 | $\chi^2=0.66, p=.416$ |
| THW (s) - Irr | 1.79±0.41 | 1.76±0.43 | 1.38±0.25 | 1.85±1.07 | $\chi^2=0.48, p=.488$ |
| THW (s) - Reg | 1.61±0.33 | 1.50±0.31 | 1.38±0.23 | 1.33±0.17 | $\chi^2=0.37, p=.545$ |

Note: Irr = Irregular, Reg = Regular

Table 3 provides a summary of descriptive and inferential statistics for the general driving task.

**Table 3.** Descriptive and inferential statistics for the **general driving task** performance by group and time point.

| Dependent Variable | Mean±CI |  |  |  | SESSION x TIME<br>interaction<br>Chi-square and p-value |
| --- | --- | --- | --- | --- | --- |
|  | tACS |  | sham |  |  |
|  | Session 0 | Session 6 | Session 0 | Session 6 |  |
| Speed variability (km/h) | 2.94±0.37 | 3.45±0.75 | 2.82±0.48 | 2.70±0.43 | $\chi^2=1.03, p=.309$ |
| Median speed (km/h) | 50.64±0.62 | 52.35±1.65 | 50.6±0.70 | 50.53±0.68 | $\chi^2=6.32, p=.012$ |
| Speeding (>5 km/h) | 8.09±4.29 | 20.19±14.41 | 7.15±5.00 | 4.04±3.02 | $\chi^2=3.29, p=.070$ |
| Lane variability (m) | 0.26±0.04 | 0.26±0.04 | 0.25±0.03 | 0.27±0.04 | $\chi^2=0.08, p=.782$ |
| DRT false alarms (N) | 1.00±0.45 | 2.15±1.18 | 1.06±0.63 | 1.58±0.79 | $\chi^2=0.54, p=.464$ |
| DRT accuracy (%) | 89.72±3.34 | 90.68±1.65 | 91.17±1.86 | 93.32±1.97 | $\chi^2=0.128, p=.721$ |
| DRT response time (ms) | 610±29 | 572±39 | 579±37 | 565±35 | $\chi^2= 4.91, p= .026$ |

Note: DRT = Detection Response Task

There was no significant interaction between SESSION and TIME for lane variability, which suggests that tACS did not affect how well participants maintained a consistent position on the road. There was a significant interaction for median speed. Follow-up contrasts revealed that median speed increased by 1.71 km/h in the tACS group between session 0 and session 6, *z* = 3.61, *p* < .001, *d* = 1.37, while the sham group exhibited no significant changes, −0.07km/h, *z* = 0.01, *p* = .996, *d* = 0.002. Importantly however, there was no corresponding increase in speeding (i.e., exceeding the limit) for the tACS group relative to the spam group. There was also no increase in speed variability, suggesting that the tACS group maintained their faster speed consistently. Given that participants were instructed to drive as close as possible to 50 km/h, the tACS group demonstrated improved task performance without a significant increase in risky behaviour (i.e., speeding). Taken together, these results suggest that tACS enhanced driving performance without compromising safety.

For the visual detection response task (DRT), there was a significant interaction between SESSION and TIME on response time, where the tACS group responded faster when detecting targets between session 0 and session 6, −38 ms, *z* = 3.91, *p* < .001, *d* = 1.502, while the sham group showed no statistically significant changes, −14 ms, *z* = 0.69, *p* = .493, *d* = 0.273. There was no interaction for DRT accuracy. This suggests that participants in the tACS group responded more quickly to DRT stimuli and may therefore have improved spare cognitive capacity while driving.

## Discussion

The current study investigated whether repeated applications (i.e., 5 sessions in 2 weeks) of bifocal tACS at beta frequency targeting the rIFG and preSMA, cortical nodes that are thought to play an important role in response inhibition enhance functional connectivity between these regions and improve response inhibition in healthy young adults. Notably, the focus on repeated applications of tACS in this research represents a novel approach, as most existing studies have examined the effects of tACS in a single session. Furthermore, we explored the potential effect of tACS-induced modulations in the brain on the real-world skill of driving performance, which requires the orchestration of multiple cognitive functions, representing a novel investigation in this field.

The EEG ImCoh measure indicated improved functional connectivity between the rIFG and preSMA following repeated tACS sessions, while no significant effects of tACS on response inhibition or braking task were observed either within or between sessions. However, for the general driving task, compared to the baseline, participants drove closer to the speed limit and showed an increased capacity for cognitive load, suggesting the potential benefit of bifocal tACS on multiple brain functions despite specifically targeting the response inhibition network. This pattern was absent in the sham group, indicating that the observed effects were specific to the active stimulation. It is important to note that the general driving task primarily reflects fundamental vehicle control, which does not directly engage response inhibition processes. While we observed changes in brain connectivity, these did not correspond to changes in SSRT or braking task performance, suggesting that the observed improvements may reflect more general enhancements in cognitive-motor integration rather than a specific effect on the response inhibition network. These findings contribute to our understanding of cumulative tACS impacts and their potential real-world applications in complex skills like driving.

### Repeated application of bifocal tACS improves functional connectivity between the rIFG and preSMA over time

In support of our previous studies (Fujiyama et al., 2023), we found no immediate change to rsEEG functional connectivity following bifocal tACS. However, the between-session analysis revealed sustained increases in functional synchrony (ImCoh) between the rIFG and preSMA, consistent with previous evidence of repeated single-site tACS sessions producing enduring effects on connectivity (Khan et al., 2023). Specifically, the tACS group showed significant increases in ImCoh from Session 1 to Session 5 in our study, which was sustained at the 7-day follow-up. This is a particularly novel finding, as to our knowledge, this is the first demonstration of significant improvements in functional connectivity following multi-session stimulation using *bifocal* tACS, specifically targeting connectivity between two cortical regions. Encouragingly, our results provide further evidence that repeated applications of bifocal tACS can lead to significant and long-lasting changes in brain connectivity, particularly within networks associated with cognitive functions.

While we found improved functional connectivity in the active tACS groups, it is important to note that the sham group exhibited a decline in ImCoh across sessions, possibly suggesting the unstable nature of ImCoh from session to session. However, as the tACS group showed the opposite pattern of change, the change in the sham group does not undermine our findings, as it shows that bifocal tACS has a distinct effect on functional connectivity when measured with ImCoh. Given that the current study used in-phase stimulation, i.e., the phase relationship between two sinusoidal currents was 0°, future research is warranted to include anti-phase (180° phase relationship) stimulation to better delineate the impact of in-phase bifocal tACS on functional connectivity.

### Repeated application of bifocal tACS did not improve response inhibition across sessions

Repeated applications of bifocal tACS at 20 Hz (i.e. beta frequency) over the rIFG and preSMA did not produce significant changes in response inhibition, nor were there improvements observed within sessions. This appeared to contrast with prior findings, which demonstrated that cognitive functions such as working memory (Reinhart & Nguyen, 2019) and response inhibition (Fujiyama et al., 2023) can be enhanced by a single session of bifocal tACS. Additionally, repeated applications of bifocal tACS have been shown to result in lasting cognitive improvements (Grover et al., 2022). The absence of significant improvements in response inhibition may reflect differences in the timing of stimulation relative to the task. Our previous study (Fujiyama et al., 2023) which compared the effects of a single session of bifocal tACS applied during versus prior to the SST, found that response inhibition improved but only in the group that received stimulation immediately prior to task performance. In the current study, we applied bifocal tACS during the task, providing further evidence that “online” bifocal tACS does not improve response inhibition when assessed with the SST.

With respect to the between-session changes, while Grover et al. (2022) found that repeated applications of bifocal tACS (4 consecutive days) resulted in lasting cognitive improvements, it is important to note that their study employed a different cohort, i.e., older adults, while our study focused on healthy young adults. The age difference between the cohorts may have contributed to the differing results. Older adults often exhibit cognitive decline (Hedden & Gabrieli, 2004) and might have more room for improving their behaviour and may be more responsive to interventions like bifocal tACS. In contrast, cognitive function in healthy young adults is likely intact, potentially making it more challenging to induce notable modulations in cognitive function within the time frame of a single or repeated tACS session. This could explain why we observed no between-section effects in response inhibition. Additionally, differences in the tasks used could play a role. Response inhibition, as measured by the SST, may require more extended time frames or more stimulations to show changes compared to functions like working memory. These factors, along with potential cohort differences, highlight the complexity of tACS effects across age groups and task types. Further research into the temporal dynamics of stimulation and function-specific responses is crucial to understanding how repeated bifocal tACS improves cognitive functions.

### Repeated application of bifocal tACS improves driving performance

We did not observe a significant impact of repeated application of 20 Hz bifocal tACS over the rIFG and preSMA on performance in the braking task, a driving task that involves the ability to adapt behaviour and inhibit prepotent responses. This finding suggests that while bifocal beta tACS may influence other aspects of cognitive or motor functions, its effects did not extend to this specific measure of braking task, which involved maintaining a safe distance while following a lead vehicle that braked periodically. In contrast, general driving performance improved after five sessions of tACS intervention. Participants in the active condition drove faster, but without a corresponding increase in speeding or speed variability relative to the sham condition. This suggests that, in line with task instructions, they were better at the primary task of travelling close to the speed limit. In the active condition, participants also responded faster to the presented DRT targets. Taken together, these data indicate that not only did participants who received tACS drive better, but they did so while retaining more spare cognitive capacity (Castro et al., 2019). In other words, driving generated less cognitive load following tACS as compared to sham, potentially leaving tACS participants with more spare capacity to respond to the DRT. Our results align with Facchin et al. (2023), who observed improved attentional capacity, measured by a detection response task following three sessions of transcranial direct current stimulation, another form of transcranial electrical stimulation (tDCS), targeting the frontal eye field (FEF), a region implicated in target selection and distractor suppression (e.g., Corbetta & Shulman, 2002). Similarly, previous tDCS studies targeting the dorsolateral prefrontal cortex (DLPFC) have shown positive effects on driving. Sakai et al. (2014) found that tDCS with an anodal electrode over the right DLPFC enhanced car-following and lane-keeping behaviour, while Beeli et al. (2008) demonstrated that tDCS with an anodal electrode over the left and right DLPFC reduced risky driving behaviour, but did not impact participants’ ability to maintain a safe distance when following a periodically braking lead vehicle.

In the current study, targeting the rIFG and preSMA with bifocal tACS demonstrated a distinct mechanism for improving general driving performance. Unlike prior tDCS studies focusing on the DLPFC and FEF, bifocal tACS applied to regions (rIFG and preSMA) that are implicated in inhibitory function resulted in a significant increase in driving speed and improved response time on a DRT. While these driving measures do not specifically assess response inhibition, the results suggest potential broader effects of tACS on cognitive and motor functions relevant to driving performance.

Taken all together, previous findings and our current results highlight the importance of exploring how stimulation of specific brain regions can improve particular functions, which would offer a foundation for tailoring interventions to target distinct cognitive and behavioural outcomes.

### Limitations and future perspectives

The present study found that functional connectivity between key regions implicated in response inhibition improved across sessions, while the behavioural manifestation of response inhibition showed no changes following bifocal tACS. The lack of significant changes in response inhibition may raise questions about the efficacy of the bifocal tACS protocol used. Individual response variability to tACS may account for this discrepancy, indicating that a one-size-fits-all approach may not be optimal for all participants. Although the tACS protocol was based on established brain oscillations linked to response inhibition (Aron et al., 2016; Swann et al., 2011), individual variability in brain oscillations could affect its effectiveness. Tailoring tACS frequencies to match participants’ unique peak oscillations might optimise performance (Veniero et al., 2019; Vosskuhl et al., 2015) or utilising a closed-loop approach to optimise stimulation parameters during the stimulation by monitoring neurophysiological readouts (Wansbrough et al., 2024) would potentially reduce individual variability. Furthermore, greater spare cognitive capacity has been related to improved driver safety and better hazard response (Strayer et al., 2015), and shown to quantify the workload induced by in-car technology distractions(Bowden et al., 2019; Strayer et al., 2017).

Our study had sufficient statistical power to detect EEG changes pre- and post-stimulation. However, future research should increase participant numbers to better investigate correlations between changes in brain connectivity and behavioural improvements. Exploratory analyses of the correlations between ImCoh ratio changes and improvements in SSRT, driving, and braking performance revealed medium effect sizes (Kendall’s Tau range: −0.6 to +0.4), but these were not statistically significant (*p*s > 0.21) and Bayesian statistics yielded anecdotal evidence (BF10s < 2.15) (Wagenmakers et al., 2018). While these findings do not establish a relationship between functional connectivity modulation and behavioural improvements, they also do not definitively rule one out. Future studies should further explore potential associations between functional connectivity changes and behavioural outcomes, particularly in the context of repeated bifocal tACS applications.

## Conclusion

The present study was the first to investigate repeated sessions of bifocal tACS over the rIFG and preSMA for improving response inhibition, functional connectivity, and driving performance. Significant improvements in functional connectivity and driving performance were observed, highlighting the potential of tACS to modulate connectivity and enhance driving performance over time. While changes in response inhibition were not detected, the extended effects on functional connectivity suggest the possibility of long-term benefits for conditions related to cortical connectivity. The lack of notable changes in response inhibition may be attributed to individual differences and parameter thresholds, as well as differences between “online” and “offline” effects of tACS. Despite this, the findings provide novel insight into the use of bifocal tACS for neuromodulation, offering a foundation for further exploration of bifocal tACS to enhance functional connectivity and associated cognitive and motor processing.

## CRediT authorship contribution statement

**Hakuei Fujiyama**: Conceptualization, Methodology, Formal analysis, Writing - Original Draft, Visualisation. **Vanessa Bowden**: Conceptualization, Methodology, Formal analysis, Writing - Review & Editing. **Alexander Tang**: Conceptualization, Methodology, Formal analysis, Writing - Review & Editing. **Jane Tan**: Conceptualization, Methodology, Investigation, Writing - Review & Editing, **Elisha Librizzi**: Investigation, Writing - Review & Editing, **Shayne Loft**: Conceptualization, Methodology, Writing - Review & Editing

## Acknowledgements

This work was supported by the Neurotrauma Research Program (DoH20193370) awarded to H.F., J.T., V.B., A.T., and S.L. A.D.T was supported by a Sarich Family Research Fellowship. The authors thank Ms Arabella Graham, Ms Tianah Linnell, Ms Alana Vincent, and Ms Paige Morgan for their contribution to the data collection.

